# Co-variation between glucocorticoids, behaviour and immunity supports the pace-of-life syndrome hypothesis: an experimental approach

**DOI:** 10.1101/2021.03.13.435238

**Authors:** Jeffrey Carbillet, Benjamin Rey, Rupert Palme, Chloé Monestier, Luca Börger, Typhaine Lavabre, Marie-Line Maublanc, Nicolas Cebe, Jean-Luc Rames, Guillaume Le Loc’h, Marine Wasniewski, Benoit Rannou, Emmanuelle Gilot-Fromont, Hélène Verheyden

## Abstract

The biomedical literature has consistently highlighted that long-term elevation of glucocorticoids might impair immune functions. In wild animals, patterns are less clear. Here, we re-explored the stress-immunity relationship considering the potential effects of behavioural profiles. Thirteen captive roe deer (*Capreolus capreolus*) were monitored over an eight-week period encompassing two capture events. We assessed how changes in baseline faecal cortisol metabolite (FCM) concentrations following a standardised capture protocol and vaccination affected changes in thirteen immune parameters of the innate and adaptive immunity, and whether behavioural profiles were linked to changes in baseline FCM levels and immune parameters. We found that individuals showing an increase in baseline FCM levels also exhibited an increase in immunity and were characterised by more reactive behavioural profiles (low activity levels, docility to manipulation and neophilia). Our results suggest that immunity of large mammals may be influenced by glucocorticoids, but also behavioural profiles, as it is predicted by the pace-of-life syndrome hypothesis. Our results highlight the need to consider co-variations between behaviour, immunity and glucocorticoids in order to improve our understanding of the among-individual variability in the stress-immunity relationships observed in wildlife, as they may be underpinned by different life-history strategies.

## Introduction

The immune system is one of the most important mechanisms in vertebrates for improving survival. This complex system is composed of two complementary arms, the innate (relatively fast and non-specific) and adaptive (slower at first encounter, but more long-lasting and specific) immunity, each composed of numerous cells and effectors [1]. This system, however, is not cost-free [1, 2], suggesting trade-offs between immune defences and other functions that use a common resource and contributes to fitness [3, 4]. Glucocorticoids (such as cortisol and corticosterone) are metabolic hormones that play a major role in the regulation of energy use [5, 6] and may therefore underlie such trade-offs.

Glucocorticoids are also one of the main mediators of the stress response. In response to external or internal stimuli, the activation of behavioural and physiological responses allows an organism to cope with challenges [7, 8]. In particular, the activation of the hypothalamic-pituitary-adrenal (HPA) axis that results in the secretion of glucocorticoids helps organisms to cope with stressful situations by making stored energy available [9]. However, repeated or chronic elevation of glucocorticoids may have negative effects on other energy-demanding functions such as reproduction [10] and immunity [11].

Over the past years, several studies investigated the relationship between stress and immunity, particularly in the biomedical domain where it has generally been shown that short-term elevation of glucocorticoids (i.e., few minutes to few hours) stimulates immune functions [12, 13], whereas chronically elevated glucocorticoid levels are immunosuppressive [9, 14,]. Focusing on long-term elevation of glucocorticoids (i.e., few days to few months), studies in wildlife have shown mixed results ranging from decreased, increased, or no change in immune functions with chronic glucocorticoid elevation [15, 16, 17, 18]. Evidence is also accumulating that glucocorticoid levels do not affect all aspects of the immune system in the same manner, such that immunoglobulin production may be impaired while other parameters (T-cell mediated, or constitutive immunity) might not be affected [19, 20].

To understand the stress-immunity relationship, little consideration has been given to the link with behavioural profiles. Close links between physiology and behaviour are expected due to the underlying energetic basis of both traits [21]. Accordingly, among-individual differences in behavioural traits are linked to their physiology, including glucocorticoid secretion and immune functions [22, 23]. For instance, in wild superb fairy-wrens (*Malurus cyaneus*), individuals exhibiting proactive behavioural traits (fast exploration of a novel environment) had the lowest level of natural antibodies [24]. Conversely, in several species, slower explorer or more reactive individuals tend to exhibit higher baseline and stress-induced glucocorticoid levels compared to faster or more proactive ones [22, 25]. In addition, a recent study on laying hens (*Gallus gallus domesticus*) highlighted that more reactive individuals exhibited greater stress and immunological (swelling in response to phytohemagglutinin injection) responsiveness than more proactive ones [26]. Such co-variations between behavioural and physiological traits can be interpreted within the pace-of-life syndrome hypothesis formulated by Réale et al. [23]. This hypothesis posits that species, populations or individuals experiencing different ecological conditions should differ in a suite of behavioural, physiological and life-history traits that may have co-evolved according to the particular ecological conditions encountered, leading to differences in life-history strategies. Within this hypothesis, individuals with slower life history strategies are expected to have more reactive behavioural profiles, higher glucocorticoid levels and higher investment in overall immunity, while those with faster life history strategies should have more proactive behavioural profiles, lower glucocorticoid levels and lower overall investment in immunity. Empirical data is however lacking to support this hypothesis.

In the present study, we investigated the relationships between changes in baseline glucocorticoid levels and changes in immunity, and how these were related to behavioural profiles. To do so, we investigated the link between variations in baseline faecal cortisol metabolite (FCM) levels and variations in thirteen adaptive and innate immune parameters, before and after a standardised capture stress protocol associated with an immune challenge using anti-rabies vaccination, in captive roe deer (*Capreolus capreolus*). In addition, we evaluated how these changes may be related to behavioural profiles, as characterised by three commonly used behavioural traits: docility, neophobia, and activity levels.

We expected that 1) changes in baseline FCM levels between the two observation periods (before/after capture) would co-vary with changes in immune parameters; 2) changes in baseline FCM levels should be less related to adaptive than innate immune parameters and inflammatory markers, due to the relatively low cost of adaptive immunity [27]. We also expected 3) that baseline FCM levels as well as variations in baseline FCM levels should be related to individual behavioural profiles [22, 28], with higher baseline levels and higher increase in baseline levels in more docile, less active and more neophilic individuals (i.e., more reactive individuals). Finally, we predicted 4) that the increase in immune parameters between the two observation periods should be greater for the most reactive individuals, which are expected to invest more in overall immunity [23, 26].

## Material and methods

### Study site

The study was conducted on a captive population of roe deer living in the Gardouch research station, located in south-west of France. The station is owned and managed by the French Research Institute for Agriculture, Food and Environment (INRAE). It consists of 12 enclosures of 0.5 ha with meadow, each containing between one to six captive roe deer, supplemented with food pellets. The experiment included 13 females, aged from 4 to 13 years old, and raised at the station from their birth or their first year of life. All had some degree of habituation to humans but were able to express normal behavioural responses (e.g., vigilance, escape) to stressful situations.

### Experimental design

The experimental procedure was carried out between mid-September and mid-November 2018 and is summarised in figure 1. During period 1, to assess baseline glucocorticoid level of each individual, we collected faeces every four days during four weeks and measured FCM concentrations. Faeces were collected immediately after defecation was observed and kept at +4°C for a maximum of 1 h before being stored at − 20°C until steroid analysis. At the end of period 1, each roe deer was subjected to a standardised capture stress protocol involving restrained immobilisation [29]. During capture, we collected faeces from the rectum, collected blood samples, injected an anti-rabies vaccine (Rabisin^®^, Merial, France, 1 ml) subcutaneously, and fitted collars equipped with tri-axial accelerometers (see Capture protocol and data collection for details). Collection of faecal samples was then continued every four days for an additional four weeks, as described above (i.e., period 2), in order to evaluate the effect of capture on baseline glucocorticoid level for each individual. Period 2 started two days after capture 1, in order to avoid measuring the acute increase in glucocorticoid level due to capture [30]. At the end of period 2, roe deer were recaptured following the same procedure (capture 2) and faeces and blood were again collected.

**Figure 1.**
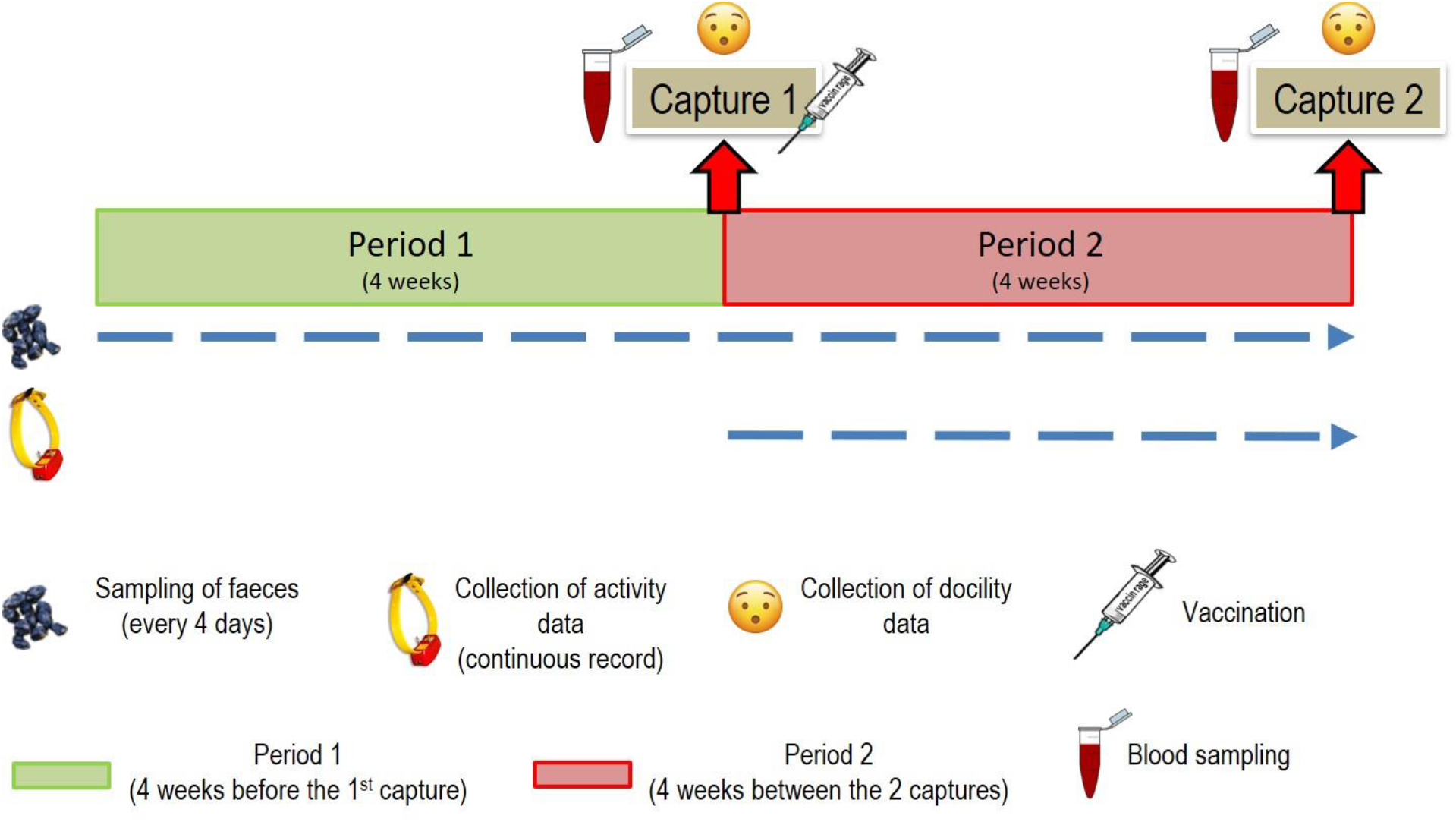
Summary of the experimental design. Data relative to the assessment of neophobia scores were obtained prior to this protocol (in February 2015 for all individuals, except two that were assessed for neophobia in February 2018, following the same protocol, see details in [40]).

### Capture protocol and data collection

Roe deer were directed into their hut by slowly approaching them and then pushed through a trap door into a retention box. Once in the box, animals were tranquilised with an intramuscular injection of acepromazine (Calmivet®, Vetoquinol, France; targeted dose of 0.075 mg/kg) [31]. Individuals were weighed with an electronic balance to the nearest 100 g.

In addition, we characterised each individual behavioural profiles using three behavioural traits, docility to capture, activity level and neophilia, at three different moments. We point out that, here, behavioural profiles do not refer to personality or behavioural syndromes, which would require repeated measures of each behavioural traits considered, and to partition phenotypic (co)variation at the among-individual versus residual levels, which was not possible to do with our data. First, docility was indexed during handling as follows: struggling (score of 1), not struggling (score of 0). This has been shown to be repeatable over time (r=0.26) with a tendency to be heritable (h=0.17) in roe deer [32]. The second trait, spontaneous daily activity [33] was measured using accelerometery data recorded at 20 Hz from tri axial accelerometers (Daily Diary tags, Wildbytes Ltd., Swansea University) mounted on animal collars. We calculated the Vectorial Dynamic Body Acceleration (VeDBA) metric [34], using a 2 second smoothing windows and the DDMT software (Wildbytes Ltd., Swansea University). VeDBA values were summed for each individual, date, and hour of the day (‘total VeDBA’) and averaged through the four weeks between the two capture events (period 2) to index daily activity. The third measured trait, neophobia, was defined as the avoidance of novel stimuli in the environment [33] and was assed using the difference in feeding efficiency with and without the presence of a novel object [35]. We calculated the ratio of the number of visits to the hut that resulted in a successful feeding bout (numerator) and the total number of visits to the hut (denominator). Measurements were repeated for five days for each condition (with and without novel object), and the difference in the ratio between the two conditions was calculated (see [35] for details). More neophobic individuals should be less inclined to feed during a given visit when a novel object is present, resulting in a higher score on the neophilia-neophobia continuum.

### Immune parameters measurement

Blood samples were taken on EDTA and dry tubes. EDTA blood was preserved at 4°C and served to measure the total leukocyte concentration (white blood cell [WBC]) with an automat (Sysmex 2000iV, Sysmex). A differential cell count (neutrophil, basophil, eosinophil, lymphocyte and monocyte) was performed on the first 100 WBCs on Wright-Giemsa-stained blood smears [36]. To obtain concentrations of each leukocyte type, the total leukocyte count was multiplied by the proportion of each cell type. The serum, was obtained after blood centrifugation (1500 g for 15 min) and was stored at -20°C for subsequent measures of total proteins, using a refractometer, albumin and alpha-1, alpha-2, beta, and gamma globulins using electrophoresis on agarose gel. Haptoglobin, an alpha-2-globulin, was also measured by spectrophotometry (Konelab 30i PLC, Fisher Thermo Scientific) Circulating levels of natural antibodies (NAbs) were measured by a hemagglutination test (HA), that measures NAbs ability to agglutinate exogenous cells, while the complement activity was revealed by the ability of proteins to induce hemolysis (HL) [37, 38]. Finally, we quantified the level of anti-rabies antibody following the method described by Cliquet et al. [39]. We therefore measured 6 markers of innate immunity (neutrophils, basophils, monocytes, eosinophils, hemagglutination and hemolysis titers), 4 inflammatory markers (haptoglobin, alpha-1, alpha-2 and beta-2 globulins), and 3 markers of adaptive immunity (lymphocytes, gamma-globulins and anti-rabies antibodies).

### Extraction and quantification of FCMs

FCMs were extracted following a methanol-based procedure and assayed using a group-specific 11-oxoaetiocholanolone enzyme immunoassay (EIA), as previously described [40] and validated for roe deer [41]. Measurements were carried out in duplicate (intra- and inter-assay coefficients of all samples were less than 10% and 15%, respectively).

### Ethical approval

All applicable institutional and/or national guidelines for the care and use of animals were followed. The protocol was approved by the Ethical Committee 115 of Toulouse and was authorized by the French government (APAFIS#14706 – 12-11-2018).

### Statistical analyses

#### Relationship between behaviour and changes in immunity and baseline FCMs

Changes in immunity (Δ immunity) were calculated for each parameter as the difference between the measurements obtained at the two capture events. Similarly, changes in baseline FCM levels (Δ glucocorticoids), were calculated as the difference of averaged baseline FCM levels between period 2 and period 1 for each individual. In addition, we used the behavioural scores at capture as an index of docility. Values did not differ between the two captures within-individual, except for one individual for which the score passed from 1 to 0. We chose to use values from the first capture in order to avoid a potential habituation effect for this individual.

Then, to test our hypotheses, we used Partial Least Square Path Modelling (PLS-PM) analysis [42]. This statistical analysis is particularly recommended when dealing with variables showing high correlation in order to avoid redundancies and high type I error [42]. Here we built the following blocks of variables, each being summarised by a latent variable: Δ glucocorticoids (1 variable), Δ innate immunity (6 variables), Δ adaptive immunity (3 variables), Δ inflammatory markers (4 variables), and behavioural profile (3 variables, electronic supplementary material table S1).

We then ran 3 PLS-PM analyses, each one evaluating the relationships between behavioural profile, change in baseline glucocorticoids, and change in 1) innate immunity, 2) adaptive immunity, and 3) inflammatory markers. For each of the three analyses, we built a structural model (or inner model, i.e., describing relationships among latent variables) that consisted of three latent variables: Δ glucocorticoids, behavioural profile, and Δ immunity (innate, adaptive or inflammatory). The statement for the structural models was as follows: change in immunity depends on both behavioural profile and change in baseline glucocorticoids, which also depends on behavioural profile. Finally, in the measurement model (or outer model, i.e., relationships between observed and latent variables), the observed variables were considered as reflecting the corresponding latent variable (reflective mode), except for innate immunity where observed variables were considered as constituting the latent variable (formative mode). This option was chosen due to the high number of biomarkers used and the complexity and diversity of the biological actions of the innate immune system [43]. This diversity is reflected in the moderate correlation among components of this latent variable.

We then ran PLS-PM analysis to adjust both the structural (figure 2) and measurement models, through multiple linear regressions. Lastly, we performed the diagnosis of each model (communality and redundancy are presented in table S2) following the recommendations of Gaston Sanchez [42]. The structural models were checked using R^2^, redundancy index (ability to predict) and Goodness-of-fit index, a pseudo GoF measure that reflects the overall prediction power of the model (0 < Gof < 1).

**Figure 2.**
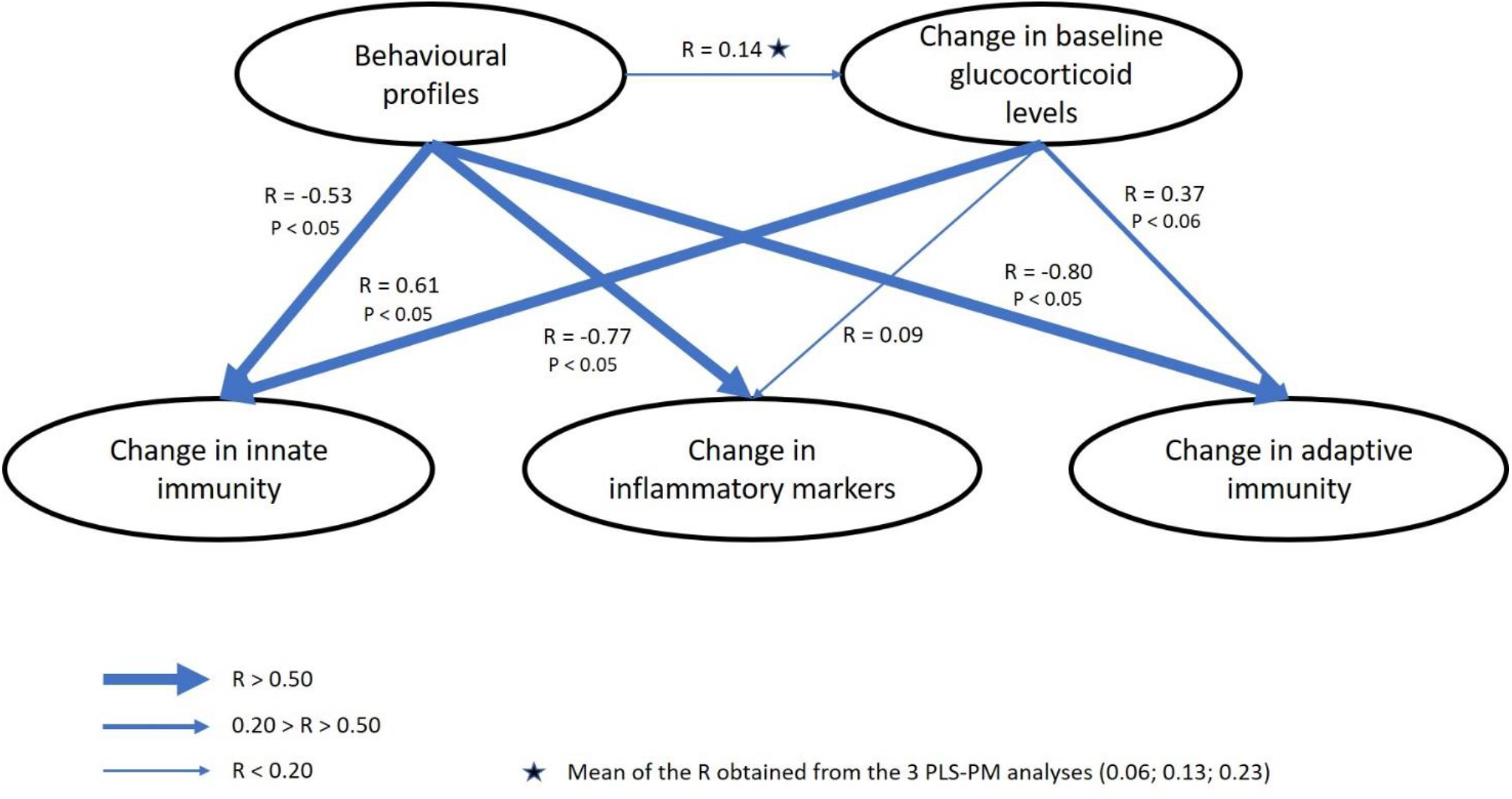
Structural models of relationships among behavioural profiles, change in baseline glucocorticoid levels and change in immunity as determined by the partial least squares path modelling analyses. Arrows indicates the direction of effect and the thickness of arrows indicates the strength of the correlation between latent variables. Absence of p-value (P) indicate the relationship was not significant.

#### Relationship between baseline FCMs throughout the experiment and behaviour

In order to test the hypothesis that baseline FCM levels throughout the experiment (period 1 + period 2) should be higher in more docile, less active and more neophilic individuals (i.e., more reactive individuals), while controlling for other factors affecting FCM levels, we performed linear mixed-effects models (LMMs) on the 181 observations of FCM levels from 13 individuals (14 repetitions per individuals with 1 missing value for one individual). FCM values were log transformed to achieve normality of model residuals. We analysed the overall correlation pattern between docility, neophobia and activity using a normed Principal Component Analysis (PCA) and used scores from the first principal component (PC1) which indexed individual proactive-reactive gradient of behaviour (electronic supplementary material table S3, figure S3). We then built a reference model that included all biologically relevant variables to explain baseline glucocorticoids levels, and compared this model with all its sub-models. The reference model included: PC1, age of individuals, and Julian date of sampling. Individual identity and enclosure identity were included as random effects to avoid pseudo-replication issues [44] and to control for unexplained variance due to among-individual differences and among-enclosure variation.

The best models of variation in FCM levels were selected based on the second-order Akaike Information Criterion (AICc) [45]. Models with a difference in AICc (ΔAICc) > 2 units from the best model were considered to have less support [45]. In addition, we removed models within two AICc units of the top model that differed from a higher-ranking model by the addition of one or more parameters, as recommended [46]. In addition, we calculated AICc weights (AICcw) to measure the relative likelihood that a given model was the best among the set of fitted models. The normality of model residuals was tested (Shapiro-Wilk test) and visually assessed. Goodness-of-fit was assessed by conditional and marginal R^2^ values and standard residual plot techniques [47].

All analyses were carried out with R version 3.6.0 [48], using the lmer function from the lme4 package [49] and the plspm function from the plspm package [42].

## Results

### Co-variation between behaviour and changes in immunity and baseline FCMs

Among the 13 individuals considered in our study, 9 showed a decrease in baseline FCM levels during period 2 compared to period 1 (ranging from -786 to -8 ng/g of wet faeces), while 4 showed an increase (ranging from 93 to 346 ng/g). In addition, for each individual, vaccination increased the level of anti-rabies antibody, but large among-individual differences were observed, with values ranging from +0.60 to +41.50 IU, with a median of +10.39. Proactive individuals were characterised by high daily activity levels, lack of docility and neophobia (table S3). In addition, the 13 individuals appeared to be homogeneously distributed along the gradient ranging from proactive to reactive behavioural profiles as showed by the PC1 axis scores ranging from -1.92 to 2.66, with a median value of 0.20 (figure S3).

Our analyses revealed links between behavioural profiles and changes in the three studied aspects of immunity (table 1). Individuals that exhibited more proactive behaviour, expressed by high daily activity levels, lack of docility and neophobia, showed an overall strong decrease in innate (r = -0.53; P < 0.05; figure 2), adaptive (r = -0.80; P < 0.001; figure 2), and inflammatory (r = -0.77; P < 0.005; figure 2) markers of immunity. However, the weights of observed variables in the definition of latent variables differed according to the analysis. When analysing Δ adaptive immunity, gamma globulins, lymphocytes and anti-rabies antibodies contributed similarly to the latent variable (weights [w] of 0.37; 0.52 and 0.54 respectively), while behavioural profile was essentially represented by docility and neophobia (w = 0.65 and 0.64 respectively). On the opposite, for Δ inflammatory markers, behavioural profile was largely represented by mean daily activity levels (w = 1.05) and less by docility (w = -0.32) and neophobia (w = 0.12), while markers of inflammation contributed overall to the same proportion to their latent variable (table 2). Lastly, for Δ innate immunity, behavioural traits contributed to the same extent to their latent variable (table 2). It is also important to note that among innate immune parameters, neutrophils were correlated negatively to other biomarkers (see the negative loading in table 2), thus the negative relationship between behavioural profiles and innate immunity only occurred for these markers, while temporal changes in neutrophil concentrations were actually positively linked to activity, neophobia and lack of docility.

**Table 1.** Characteristics of the partial least squares path modelling analyses to explain the relationships between behavioural profiles, change in baseline glucocorticoid levels, and change in innate, adaptive and inflammatory markers of immunity. GoF indicates the goodness of fit of the model. SE stands for Standard Error. See text for definition of the observed variables that composed each latent variable. Variables in bold represent endogenous variables.

| Parameter | Estimate | SE | t value | P value |
| --- | --- | --- | --- | --- |
| Structural model for innate immunity biomarkers (GoF: 0.32) |  |  |  |  |
| **FCMs** |  |  |  |  |
| Intercept | 2.4 e-17 | 0.301 | 7.98 e-17 | 1.00 |
| Behavioural profile | 0.055 | 0.301 | 0.183 | 0.86 |
| **Innate immunity** |  |  |  |  |
| Intercept | -4.56 e-17 | 0.197 | -2.32 e-16 | 1.00 |
| Behavioural profile | -0.53 | 0.197 | -2.70 | 0.02 |
| FCMs | 0.61 | 0.197 | 3.08 | 0.01 |
| Structural model for adaptive immunity biomarkers (GoF: 0.42) |  |  |  |  |
| **FCMs** |  |  |  |  |
| Intercept | 1.15 e-17 | 0.299 | 3.84 e-17 | 1.00 |
| Behavioural profile | 0.127 | 0.299 | 0.424 | 0.68 |
| **Adaptive immunity** |  |  |  |  |
| Intercept | 3.43 e-17 | 0.172 | 2.00 e-16 | 1.00 |
| Behavioural profile | -0.801 | 0.173 | -4.63 | 0.001 |
| FCMs | 0.373 | 0.173 | 2.16 | 0.057 |
| Structural model for inflammatory biomarkers (GoF: 0.39) |  |  |  |  |
| **FCMs** |  |  |  |  |
| Intercept | 1.41 e-16 | 0.294 | 4.79 e-16 | 1.00 |
| Behavioural profile | 0.226 | 0.294 | 0.183 | 0.46 |
| **Inflammatory markers** |  |  |  |  |
| Intercept | 3.64 e-16 | 0.192 | 1.90 e-15 | 1.00 |
| Behavioural profile | -0.770 | 0.197 | -3.91 | 0.003 |
| FCMs | 0.089 | 0.197 | 0.454 | 0.660 |

**Table 2.** Characteristics of the observed variables that composed each latent variable in the three partial least squares path modelling analyses to explain the relationships between behavioural profiles, change in baseline glucocorticoid levels, and change in innate, adaptive and inflammatory immunity. Weight represents the contribution of the variable to the latent variable, and loadings indicate the direction of the correlation between the observed variables and their latent variable. Communality indicates the amount of variability in an observed variable that is captured by its latent variable, and particularly apply for reflective latent variables. Redundancy indicates the ability to predict for a given observed variable, and particularly apply for formative latent variables. See text for definition of the observed variables composing each latent variable. Variables in bold represent endogenous variables.

| Parameter | Weight | Loading |
| --- | --- | --- |
| Outer model for innate immunity biomarkers |  |  |
| **Behavioural profile** |  |  |
| Lack of docility | 0.521 | 0.847 |
| Neophobia | 0.434 | 0.753 |
| Activity | 0.358 | 0.648 |
| **Glucocorticoids** |  |  |
| FCMs | 1.00 | 1.00 |
| **Innate immunity** |  |  |
| Neutrophils | -0.324 | -0.686 |
| Eosinophils | 0.396 | 0.495 |
| Monocytes | -0.533 | 0.274 |
| Basophils | 0.843 | 0.416 |
| Hemagglutination | 0.662 | 0.435 |
| Hemolysis | 0.245 | 0.367 |
| Outer model for adaptive immunity biomarkers |  |  |
| **Behavioural profile** |  |  |
| Lack of docility | 0.551 | 0.824 |
| Neophobia | 0.637 | 0.877 |
| Activity | -0.041 | 0.306 |
| **Glucocorticoids** |  |  |
| FCMs | 1.00 | 1.00 |
| **Adaptive immunity** |  |  |
| Gamma globulins | 0.365 | 0.581 |
| Lymphocytes | 0.523 | 0.754 |
| Anti-rabies antibody | 0.543 | 0.725 |
| Outer model for inflammatory immunity biomarkers |  |  |
| **Behavioural profile** |  |  |
| Lack of docility | -0.319 | 0.115 |
| Neophobia | 0.115 | 0.215 |
| Activity | 1.051 | 0.962 |
| **Glucocorticoids** |  |  |
| FCMs | 1.00 | 1.00 |
| **Inflammatory immunity** |  |  |
| Alpha-1 globulins | 0.360 | 0.936 |
| Alpha-2 globulins | 0.279 | 0.618 |
| Beta-globulins | 0.217 | 0.456 |
| Haptoglobin | 0.456 | 0.859 |

Changes in baseline glucocorticoid levels were associated with changes in innate (r = 0.61; P < 0.05) and adaptive immunity (tendency: r = 0.37; P < 0.06), but not inflammatory markers (table 1; figure 2). Individuals that underwent an increase in baseline FCM levels between periods also exhibited an increase in both innate and adaptive immunity. However, as pointed out above, this positive relationship means that individuals exhibiting an increase in baseline FCM levels actually had a decrease in neutrophil concentration.

Finally, the relationship between behavioural profile and change in baseline FCM levels was non-significant for all three models (table 1).

### Co-variation between baseline FCM levels throughout the experiment and behaviour

According to the model selection procedure, the best model describing among-individual differences in baseline FCMs throughout the experiment in relation to individual behavioural profiles included PC1 score and period of the experimental protocol (table S4). Specifically, roe deer that exhibited a more reactive behavioural profile (low daily activity levels, docility and neophilia) also exhibited higher baseline FCM levels throughout the experiment compared to roe deer exhibiting a more proactive behavioural profile (0.103; P < 0.005; table 3; figure 3). In addition, baseline FCM levels decreased during the second part of the experimental protocol compared to the first one (−0.1663; P < 0.05; table 3), meaning that the mean baseline FCM level of the studied roe deer population was lower during period 2 than during period 1. Age and Julian date did not increase the fit of the model.

**Table 3.** Characteristics of the selected linear mixed-effect models for explaining variation in baseline FCM levels (log-transformed) in the roe deer population of Gardouch. The effect of PC1 (behavioural profile ranging from proactive behavioural profiles to reactive behavioural profiles), age of individuals, period of sample collection, and Julian date of sample collection were fitted. Models included individual identity and enclosure number as random effects. R^2m^ and R^2c^ are the marginal and conditional explained variance of the models, respectively. SE stands for Standard Error. See text for definition of model sets.

| Parameter | Estimate | SE | t-value | P-value |
| --- | --- | --- | --- | --- |
| ($R^{2m}$: 0.06 ; $R^{2c}$: 0.27) | | | | |
| Intercept | 6.638 | 0.141 | 47.10 | < 0.001 |
| Behavioural profile (PC1) | 0.103 | 0.036 | 2.849 | < 0.005 |
| Period (2) | -0.163 | 0.080 | -2.045 | < 0.05 |

**Figure 3.**
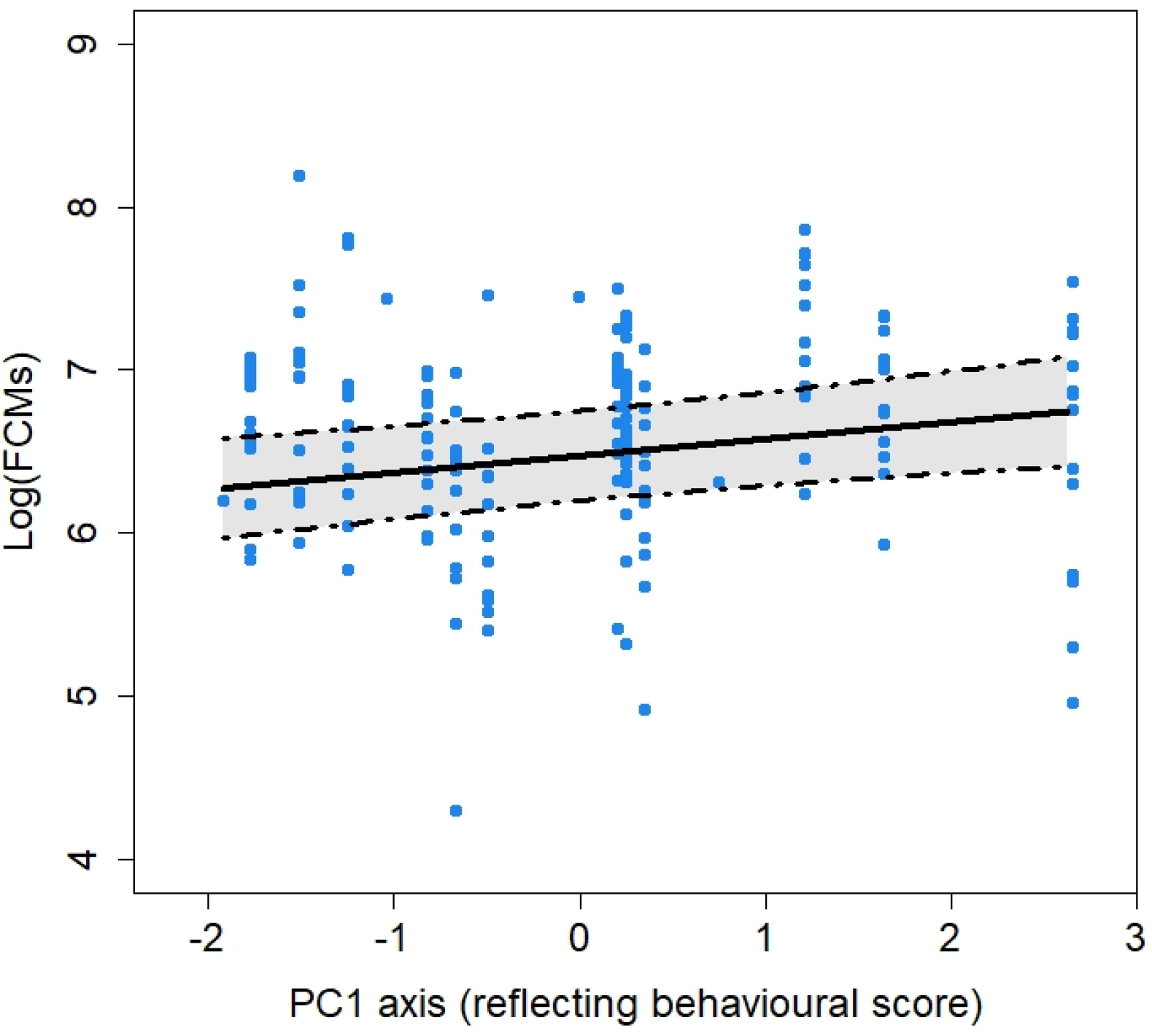
Relationship between baseline FCMs level (log-transformed) and behavioural profiles. Behavioural profiles’ scores correspond to the score for the first axis (PC1) of the PCA conducted using docility, activity and neophilia as co-variables. The three variables were all negatively correlated with PC1. Thus, this axis represents a gradient of behavioural profiles, with negative values indicating proactive behavioural profiles (high activity levels, neophobia, and lack of docility), and positive values indicating reactive behavioural profiles (low activity levels, neophilia, and docility). Points represent observed values, lines represent model predictions and dashed lines represent the 95% confidence interval.

## Discussion

In this study, we used an experimental approach to gain a better understanding on how changes in baseline glucocorticoid levels may affect simultaneous changes in immune parameters of the innate, adaptive and inflammatory markers of immunity, at the scale of few weeks (8 weeks). Our results demonstrated that an increase in baseline FCM levels was associated with an increase in immune parameters of the innate and adaptive arms, but not in inflammation. Secondly, we tested whether behavioural profiles could influence the co-variation between changes in immune parameters and baseline glucocorticoid levels. As predicted, behavioural profiles appeared to be strongly linked to changes in overall immunity, but also to baseline glucocorticoid levels throughout the experiment, while there were not related to changes in baseline glucocorticoids between the two periods of the study.

An increase in baseline glucocorticoid levels between the two periods of the study was generally related to an increase in innate immunity, except for neutrophil concentrations, which decreased as glucocorticoid levels increased. The observed negative relationship with neutrophils is consistent with the previous finding of an immunosuppressive effect of long-term elevation of glucocorticoids on immunity [11, 13]. However, the overall increase in innate immunity was unexpected and contrary to our predictions. One of the assumptions underlying the hypothesis of a negative relationship between immunity and glucocorticoids is the energy cost of immunity, which should subsequently trade-off against other energy demanding functions [2]. However, it is important to note that in our studied population, resources are not limiting and roe deer are not exposed to unpredictable variations in food resources. This particular context may diminish our ability to detect trade-offs between energy demanding functions. Alternatively, it has been proposed that, as the main function of the stress response is to recover from stressors, a decrease in immunity should not necessarily occur when glucocorticoids increase, as it could improve survival [11].

While the above hypothesis may partly explain the link between change in innate immunity and change in baseline glucocorticoids, it does not explain the difference observed between neutrophils and other biomarkers of innate immunity. Neutrophils are part of the cellular immunity and reflect acute inflammatory response while monocytes reflect chronic inflammatory response, and hemagglutination and hemolysis are both part of the humoral innate response [43]. Basophils are particularly secreted in presence of ticks [50] that are frequently encountered in the experimental facility (unpublished data). Finally, eosinophils are known to specifically bridge innate and adaptive immunity [43]. Considering the differences in the functions of these biomarkers, it is likely that they are not all linked to glucocorticoids in the same way. This could explain that we did not detect any trade-off, and that positive relationship between changes in glucocorticoid levels and innate immunity may occur (e.g., for basophils in the presence of ticks).

With respect to a change in the adaptive arm of the immune system, our results did not support the finding of the immunosuppressive effect of long-term elevation of glucocorticoids, but instead showed that adaptive immune parameters increased in individuals that showed an increase in baseline FCM levels. A first possible explanation of this result could be linked to the transient increase in glucocorticoids that occurred during the first capture, where vaccination was done. Previous work showed that a short-term elevation in glucocorticoids at the time of vaccination reinforces the efficacy of vaccination [13]. A similar scenario may have occurred in our study, and the stress of capture, resulting in a sudden increase in circulating glucocorticoids, would have strengthened the immune response to anti-rabies vaccine. This would explain our result, at least in part, if there is a correlation between the change observed in baseline FCM levels and the stress-induced increase in glucocorticoids. Another possible explanation is that the energy cost of mounting an antibody response is too low [27] for a trade-off between antibody expression and other functions to be detectable. The positive relationship we observed in our study also supports the pace-of-life syndrome hypothesis, according to which individuals with higher baseline glucocorticoid levels should show stronger investment in immunity than those with lower baseline glucocorticoid levels [23].

Overall, individuals that underwent a decrease in baseline glucocorticoids may have switched their investment away from the immune system, possibly toward another energy demanding function. We suggest that such a switch could be underpinned by a plastic response to the stressful event of capture, leading to an adjustment of the individuals’ life history strategies and change in investment between functions supporting either long-term survival or current reproduction. This would be in accordance with the pace-of-life syndrome hypothesis, where a positive association is expected between glucocorticoid levels and immunity, with higher levels in individuals favouring their long-term survival, while lower levels are expected in individuals favouring reproduction and growth [23].

Regarding the association between behavioural profiles and change in immunity, we found an overall similar association for innate, adaptive and inflammatory immunity. Precisely, individuals showing a propensity to be active, non-docile to manipulation and neophobic showed a decreased investment in their immune system following the first capture event. This result is consistent with our predictions and supports the hypothesis that fast-living individuals should have a proactive behavioural profile and a low investment in immune functions that would allow them to favour immediate reproduction over survival [23]. On the opposite, individuals showing more reactive behavioural profiles are supposed to have a slower pace-of-life and are expected to favour functions enhancing survival and future reproduction [23]. Stronger investment in immunity is thus expected for these individuals as they are more likely to face repeated encounter with the same pathogens.

Finally, we investigated whether behavioural profiles could be associated with among-individual variations in baseline glucocorticoid levels throughout the experiment, and also with changes in baseline glucocorticoid levels over an 8-week period. Our results did not provide any support for the latter, but instead supported the former. Specifically, individuals exhibiting proactive behavioural profiles also showed lower baseline glucocorticoid levels throughout the experiment compared to individuals exhibiting more reactive behavioural profiles. This result supports previous studies on wild [51] and captive [29] roe deer. It is also in accordance with the coping style framework [22] and the pace-of-life syndrome hypothesis [23], which states that behavioural and physiological responses to stressful situations are correlated. The difference we observed in our results may thus suggest that activity (reflected by baseline levels) and reactivity (reflected by the changes following the first capture event) of the HPA axis may not be associated in the same manner with among-individual variations in behaviour [22].

Overall, individual roe deer responded differently to our protocol, with some individuals showing an increase in baseline glucocorticoid levels, while others showed a decrease. In addition, our results suggest that increased baseline glucocorticoid levels are associated with a re-allocation of energy resources to innate and adaptive immunity in individuals with more reactive behaviours (see figure 4 for a summary of the outcomes of the protocol). We suggest that the observed association between immunity and baseline glucocorticoid levels is associated with different life-history strategies and underpinned by energetic trade-offs between functions enhancing survival, reproduction and growth, which would be congruent with the pace-of-life syndrome hypothesis [23].

**Figure 4.**
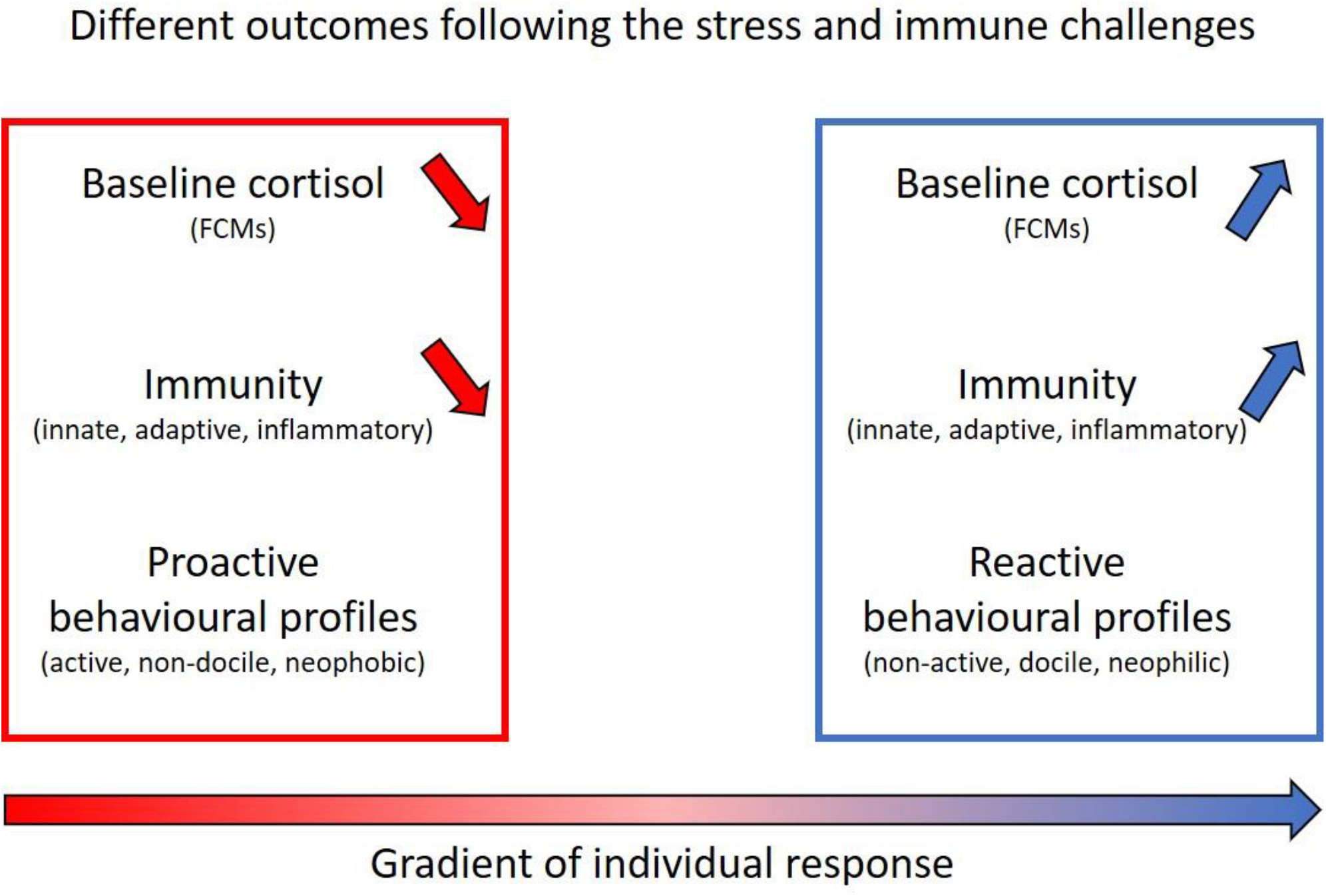
Summary of the observed outcomes and relationships between the latent variables considered in our study. Our protocol (see details in the main text) resulted in different outcomes, with part of the individuals showing an increase in baseline cortisol between the two study periods (indicated by an increase in FCMs), while others showed a decrease. Individuals that showed more reactive behavioural profiles (indicated by low activity levels, docility to manipulation, and neophilia) also exhibited an increase in baseline cortisol levels, and an increase in immunity (both innate and adaptive immunity), while the opposite occurred for individuals that showed more proactive behavioural profile.

Finally, considering that all of our results tend to support a co-variation between stress hormones, immunity and behaviour, we recommend that future work goes one step further and investigates how among-individual variations in behaviour could modulate the variation of glucocorticoid levels, as well as the relationship between glucocorticoid hormones and immunity. The extent to which co-variations between these traits are influenced by different life history strategies in the wild also warrants further investigations as it may help to understand the large and unexplained among-individual variability in the glucocorticoids-immunity relationship observed in wildlife studies.

## Supporting information

ESM S1, S2, S3, and S4

## Acknowledgments

We thank Eric Bideau and Benoit Coracin for their help throughout this study regarding roe deer manipulation and monitoring. We also thank Edith Klobetz-Rassam for EIA analysis, veterinary students for their assistance during capture events, as well as Laura Gervais for constructive discussions and comments that helped to improve this manuscript. We are also very grateful to the ECODIV department of the INRAE for its financial support to the Gardouch experimental station.

## Author contributions

JC, BRey, EGF and HV conceived and designed the study. JC, HV, MLB, NC, JLR, GLL undertook the fieldwork. LB performed the VeDBA analysis. RP performed the FCMs analysis. MW quantified anti-rabies circulating antibodies. TL, BR and EGF performed the immunological analysis. CM designed and performed the neophobia analysis. JC performed the statistical analysis, wrote the first draft of the paper and then received input from all other co-authors. All authors approved the final version of the manuscript and agree to be held accountable for the content therein.

## Data accessibility

All data and code used in this analysis are available as electronic supplementary material S5.

## References

[1] Demas GE. 2004 The energetics of immunity: a neuroendocrine link between energy balance and immune function. Horm. Behav. 45, 173–180. (https://doi.org/10.1016/j.yhbeh.2003.11.002)

[2] Lochmiller RL, Deerenberg C. 2000 Trade-offs in evolutionary immunology: just what is the cost of immunity?. Oikos 88, 87–98. (https://doi.org/10.1034/j.1600-0706.2000.880110.x)

[3] Sheldon BC, Verhulst S. 1996 Ecological immunology: costly parasite defences and trade-offs in evolutionary ecology. Trends Ecol. Evol. 11, 317–321. (https://doi.org/10.1016/0169-5347(96)10039-2)

[4] Rauw WM. 2012 Immune response from a resource allocation perspective. Front. Genet. 3, 267. (https://doi.org/10.3389/fgene.2012.00267)

[5] Landys MM, Ramenofsky M, Wingfield JC. 2006 Actions of glucocorticoids at a seasonal baseline as compared to stress-related levels in the regulation of periodic life processes. Gen. Comp. Endocrinol. 148, 132–149. (https://doi.org/10.1016/j.ygcen.2006.02.013)

[6] Busch DS, Hayward LS. 2009 Stress in a conservation context: a discussion of glucocorticoid actions and how levels change with conservation-relevant variables. Biol. Conserv. 142, 2844–2853. (https://doi.org/10.1016/j.biocon.2009.08.013)

[7] Wingfield JC, Maney DL, Breuner CW, Jacobs JD, Lynn S, Ramenofsky M, Richardson RD. 1998 Ecological bases of hormone—behavior interactions: the “emergency life history stage”. Am. Zool. 38, 191–206. (https://doi.org/10.1093/icb/38.1.191)

[8] Romero LM, Wingfield JC. 2015 Tempests, poxes, predators, and people: stress in wild animals and how they cope. Oxford University Press. (https://doi.org/10.1093/acprof:oso/9780195366693.001.0001)

[9] Sapolsky RM, Romero LM, Munck AU. 2000 How do glucocorticoids influence stress responses? Integrating permissive, suppressive, stimulatory, and preparative actions. Endocr. rev. 21, 55–89. (https://doi.org/10.1210/edrv.21.1.0389)

[10] Boonstra R. 2005. Equipped for life: the adaptive role of the stress axis in male mammals. J. Mammal. 86, 236–247. (https://doi.org/10.1644/BHE-001.1)

[11] Martin LB. 2009 Stress and immunity in wild vertebrates: timing is everything. Gen. Comp. Endocrinol. 163, 70–76. (https://doi.org/10.1016/j.ygcen.2009.03.008)

[12] Bilbo SD, Dhabhar FS, Viswanathan K, Saul A, Yellon SM, Nelson RJ. 2002 Short day lengths augment stress-induced leukocyte trafficking and stress-induced enhancement of skin immune function. Proc. Natl. Acad. Sci. U.S.A 99, 4067–4072. (https://doi.org/10.1073/pnas.062001899)

[13] Dhabhar FS. 2014 Effects of stress on immune function: the good, the bad, and the beautiful. Immunol. Res. 58, 193–210. (https://doi.org/10.1007/s12026-014-8517-0)

[14] Dhabhar FS, McEwe BS. 1997 Acute stress enhances while chronic stress suppresses cell-mediated immunity in vivo: A potential role for leukocyte trafficking. Brain Behav. Immun. 11, 286–306. (https://doi.org/10.1006/brbi.1997.0508)

[15] Matson KD, Tieleman BI, Klasing KC. 2006 Capture stress and the bactericidal competence of blood and plasma in five species of tropical birds. Physiol. Biochem. Zool. 79, 556–564. (https://doi.org/10.1086/501057)

[16] Delehanty B, Boonstra R. 2009 Impact of live trapping on stress profiles of Richardson’s ground squirrel (Spermophilus richardsonii). Gen. Comp. Endocrinol. 160, 176–182. (https://doi.org/10.1016/j.ygcen.2008.11.011)

[17] Merrill L, Angelier F, O’Loghlen AL, Rothstein SI, Wingfield JC. 2012 Sex-specific variation in brown-headed cowbird immunity following acute stress: a mechanistic approach. Oecologia 170, 25–38. (https://doi.org/10.1007/s00442-012-2281-4)

[18] Brooks KC, Mateo JM. 2013 Chronically raised glucocorticoids reduce innate immune function in Belding’s ground squirrels (Urocitellus beldingi) after an immune challenge. Gen. Comp. Endocrinol. 193, 149–157. (https://doi.org/10.1016/j.ygcen.2013.07.019)

[19] Bourgeon S, Raclot T. 2006 Corticosterone selectively decreases humoral immunity in female eiders during incubation. J. Exp. Biol. 209, 4957–4965. (https://doi.org/10.1242/jeb.02610)

[20] Stier KS, Almasi B, Gasparini J, Piault R, Roulin A, Jenni L. 2009 Effects of corticosterone on innate and humoral immune functions and oxidative stress in barn owl nestlings. J. Exp. Biol. 212, 2085–2091. (https://doi.org/10.1242/jeb.024406)

[21] Careau V, Thomas D, Humphries MM, Réale D. 2008 Energy metabolism and animal personality. Oikos 117, 641–653. (https://doi.org/10.1111/j.0030-1299.2008.16513.x)

[22] Koolhaas JM, Korte SM, De Boer SF, Van Der Vegt BJ, Van Reenen CG, Hopster H, et al. 1999 Coping styles in animals: current status in behavior and stress-physiology. Neurosci. Biobehav. Rev. 23, 925–935. (https://doi.org/10.1016/S0149-7634(99)00026-3)

[23] Réale D, Garant D, Humphries MM, Bergeron P, Careau V, Montiglio PO. 2010 Personality and the emergence of the pace-of-life syndrome concept at the population level. Philos. Trans. R. Soc. Lond, B Biol. Sci 365, 4051–4063. (https://doi.org/10.1098/rstb.2010.0208)

[24] Jacques-Hamilton R, Hall ML, Buttemer WA, Matson KD, da Silva AG, Mulder RA, Peters A. 2017 Personality and innate immune defenses in a wild bird: Evidence for the pace-of-life hypothesis. Horm. Behav. 88, 31–40. (https://doi.org/10.1016/j.yhbeh.2016.09.005)

[25] Stöwe M, Rosivall B, Drent PJ, Möstl E. 2010 Selection for fast and slow exploration affects baseline and stress-induced corticosterone excretion in Great tit nestlings, Parus major. Horm. Behav. 58, 864–871. (https://doi.org/10.1016/j.yhbeh.2010.08.011)

[26] Pusch EA, Navara KJ. 2018 Behavioral phenotype relates to physiological differences in immunological and stress responsiveness in reactive and proactive birds. Gen. Comp. Endocrinol. 261, 81–88. (https://doi.org/10.1016/j.ygcen.2018.01.027)

[27] McDade TW, Georgiev AV, Kuzawa CW. 2016 Trade-offs between acquired and innate immune defenses in humans. Evol. Med. Public Health 2016, 1–16. (https://doi.org/10.1093/emph/eov033)

[28] Montiglio PO, Garant D, Pelletier F, Réale D. 2012 Personality differences are related to long-term stress reactivity in a population of wild eastern chipmunks, Tamias striatus. Anim. Behav. 84, 1071–1079. (https://doi.org/10.1016/j.anbehav.2012.08.010)

[29] Monestier C, Gilot-Fromont E, Morellet N, Debeffe L, Cebe N, Merlet J, et al. 2016 Individual variation in an acute stress response reflects divergent coping strategies in a large herbivore. Behav. Processes 132, 22–28. (https://doi.org/10.1016/j.beproc.2016.09.004)

[30] Dehnhard M, Clauss M, Lechner-Doll M, Meyer HHD, Palme R. 2001 Noninvasive monitoring of adrenocortical activity in roe deer (Capreolus capreolus) by measurement of fecal cortisol metabolites. Gen. Comp. Endocrinol. 123, 111–120. (https://doi.org/10.1006/gcen.2001.7656)

[31] Montané J, Marco I, López-Olvera J, Perpinan D, Manteca X, Lavin S. 2003 Effects of acepromazine on capture stress in roe deer (Capreolus capreolus). J. Wild. Dis. 39, 375–386. (https://doi.org/10.7589/0090-3558-39.2.375)

[32] Gervais L, Hewison AJ, Morellet N, Bernard M, Merlet J, Cargnelutti B, Chaval Y, Pujol B, Quéméré E. 2020 Pedigree-free quantitative genetic approach provides evidence for heritability of movement tactics in wild roe deer. J. Evol. Biol. 33, 595–607. (https://doi.org/10.1111/jeb.13594)

[33] Réale D, Reader SM, Sol D, McDougall PT, Dingemanse NJ. 2007 Integrating animal temperament within ecology and evolution. Biol. Rev. 82, 291–318. (https://doi.org/10.1111/j.1469-185X.2007.00010.x)

[34] Wilson RP, Börger L, Holton MD, et al. 2020 Estimates for energy expenditure in free-living animals using acceleration proxies: A reappraisal. J. Anim. Ecol. 89, 161– 172. https://doi.org/10.1111/1365-2656.13040 (https://doi.org/10.1111/1365-2656.13040)

[35] Monestier C, Morellet N, Verheyden H, Gaillard JM, Bideau E, Denailhac A, Lourtet B, Cebe N, Picot D, Rames JM, Hewison AM. 2017 Neophobia is linked to behavioural and haematological indicators of stress in captive roe deer. Anim. Behav. 126, 135–143. (https://doi.org/10.1016/j.anbehav.2017.01.019)

[36] Houwen B. 2001 The differential cell count. Lab. Hematol. 7, 89–100.

[37] Matson KD, Ricklefs RE, Klasing KC. 2005 A hemolysis–hemagglutination assay for characterizing constitutive innate humoral immunity in wild and domestic birds. Dev. Comp. Immunol. 29, 275–286. (https://doi.org/10.1016/j.dci.2004.07.006)

[38] Gilot-Fromont E, Jégo M, Bonenfant C, Gibert P, Rannou B, Klein F, Gaillard JM. 2012 Immune phenotype and body condition in roe deer: individuals with high body condition have different, not stronger immunity. PLoS One 7, e45576. (https://doi.org/10.1371/journal.pone.0045576)

[39] Cliquet F, Aubert M, Sagne L. 1998 Development of a fluorescent antibody virus neutralisation test (FAVN test) for the quantitation of rabies-neutralising antibody. J. Immunol. Methods 212, 79–87. (https://doi.org/10.1016/S0022-1759(97)00212-3)

[40] Möstl E, Maggs JL, Schrötter G, Besenfelder U, Palme R. 2002 Measurement of cortisol metabolites in faeces of ruminants. Vet. Res. Commun. 26, 127–139. (https://doi.org/10.1023/A:1014095618125)

[41] Zbyryt A, Bubnicki JW, Kuijper DP, Dehnhard M, Churski M, Schmidt K. 2017 Do wild ungulates experience higher stress with humans than with large carnivores? Behav. Ecol. 29, 19–30. (https://doi.org/10.1093/beheco/arx142)

[42] Sanchez G. 2013 PLS path modeling with R. Berkeley: Trowchez Editions 383, 2013. (http://www.gastonsanchez.com/PLS)

[43] Demas GE, Zysling DA, Beechler BR, Muehlenbein MP, French SS. 2011 Beyond phytohaemagglutinin: assessing vertebrate immune function across ecological contexts. J. Anim. Ecol. 80, 710–730. (https://doi.org/10.1111/j.1365-2656.2011.01813.x)

[44] Hurlbert SH. 1984 Pseudoreplication and the design of ecological field experiments. Ecol. Monogr. 54, 187–211. (https://doi.org/10.2307/1942661)

[45] Burnham KP, Anderson DR. 2002 Model selection and multimodel inference: a practical information-theoretic approach. Springer Science & Business Media. (https://doi.org/10.1007/b97636)

[46] Arnold TW. 2010 Uninformative parameters and model selection using Akaike’s Information Criterion. J. Wildl. Manage. 74, 1175–1178. (https://doi.org/10.1111/j.1937-2817.2010.tb01236.x)

[47] Nakagawa S, Schielzeth H. 2013 A general and simple method for obtaining R2 from generalized linear mixed-effects models. Methods Ecol. Evol. 4, 133–142. (https://doi.org/10.1111/j.2041-210x.2012.00261.x)

[48] R Core Team (2016). R: A language and environment for statistical computing. R Foundation for Statistical Computing, Vienna, Austria. (http://www.R-project.org)

[49] Bates D, Mächler M, Bolker B, Walker S. 2014 Fitting linear mixed-effects models using lme4. arXiv preprint arXiv:1406.5823. (https://arxiv.org/abs/1406.5823v1)

[50] Karasuyama H, Tabakawa Y, Ohta T, Wada T, Yoshikawa S. 2018 Crucial Role for Basophils in Acquired Protective Immunity to Tick Infestation. Front. Physiol. 9, 1–8. (https://doi.org/10.3389/fphys.2018.01769)

[51] Carbillet J, Rey B, Lavabre T, Chaval Y, Merlet J, Débias F, Régis C, Pardonnet S, Duhayer J, Gaillard JM, et al. 2019 The Neutrophil to Lymphocyte ratio and individual coping styles across variable environments in the wild. Behav. Ecol. Sociobiol. 73, 1–13. (https://doi.org/10.1007/s00265-019-2755-z)

