## Supplementary material for "Co-variation between glucocorticoids, behaviour and immunity supports the pace-of-life syndrome hypothesis: an experimental approach": ESM S1, S2, S3, and S4: S1_ProcB_Carbillet.docx

^7 Equipe de Biologie médicale-Histologie, CREFRE, Inserm-UPS-ENVT, Toulouse, France^

^8 Inovie Vet, Laboratoire d'Analyses et Biologie Vétérinaires, Montpellier, France^

^9 Université de Toulouse, ENVT, INRAE, UMR IHAP, Toulouse, France.^

^10 ANSES, Nancy Laboratory for Rabies and Wildlife, Malzéville, France^

**Table S1.** **Summary of the latent variables used in the present study.** Observed variables are presented in blocks reflecting their associated latent variable.

| **Latent variables** | **Observed variables** | **Average values (min ; max)** |
| --- | --- | --- |
| Δ baseline glucocorticoids | Δ FCMs (ng/g of wet faeces) | -146 (-786 ; 346) |
| Δ innate immunity | Δ Neutrophils (10^3 cells/mL) | -1.23 (-4.03 ; 1.5) |
|  | Δ Eosinophils (10^3 cells/mL) | 0.01 (-0.29 ; 0.38) |
|  | Δ Basophils (10^3 cells/mL) | 0.06 (-0.36 ; 0.46) |
|  | Δ Monocytes (10^3 cells/mL) | -0.05 (-0.37 ; 0.09) |
|  | Δ Hemagglutination (titer) | -0.04 (-1 ; 1.5) |
|  | Δ Hemolysis (titer) | -0.07 (-1 ; 1.5) |
| Δ adaptive immunity | Δ Lymphocytes (10^3 cells/mL) | 0.76 (-0.80 ; 2.23) |
|  | Δ Anti-rabies antibody (IU) | 12.30 (0.60 ; 41.5) |
|  | Δ Gamma globulins (mg/mL) | 0.10 (-4 ; 3.50) |
| Δ inflammatory markers | Δ Alpha-1 globulins (mg/mL) | 0.10 (-0.20 ; 0.80) |
|  | Δ Alpha-2 globulins (mg/mL) | -0.06 (-0.90 ; 2.90) |
|  | Δ Beta-2 globulins (mg/mL) | 0.13 (-0.40 ; 0.50) |
|  | Δ Haptoglobin (mg/mL) | 0.13 (-0.26 ; 1.19) |
| Behavioural profiles | Docility score | 0.5 (0 ; 1) |
|  | Neophobia score | 0.20 (0.01 ; 0.40) |
|  | Mean daily activity | 228.8 (183.6 ; 259.9) |
