## Supplementary material for "Co-variation between glucocorticoids, behaviour and immunity supports the pace-of-life syndrome hypothesis: an experimental approach": ESM S1, S2, S3, and S4: S2_ProcB_Carbillet.docx

^7 Equipe de Biologie médicale-Histologie, CREFRE, Inserm-UPS-ENVT, Toulouse, France^

^8 Inovie Vet, Laboratoire d'Analyses et Biologie Vétérinaires, Montpellier, France^

^9 Université de Toulouse, ENVT, INRAE, UMR IHAP, Toulouse, France.^

^10 ANSES, Nancy Laboratory for Rabies and Wildlife, Malzéville, France^

**Table S2.** Additional characteristics of the observed variables that composed each latent variable in the three partial least squares path modelling analyses for explaining the relationships between behavioural profiles, change in baseline glucocorticoid levels, and change in innate, adaptive and inflammatory immunity. Communality indicates the amount of variability in an observed variable that is captured by its latent variable, and particularly apply for reflective latent variables. Redundancy indicates the ability to predict for a given observed variable, and particularly apply for formative latent variables. See text of the main document for definition of the observed variables that composed each latent variable. Variables in bold represent endogenous variables.

| **Parameter** | **Communality** | **Redundancy** |
| --- | --- | --- |
| Outer model for innate immunity biomarkers | | |
| **Behavioural profile** |  |  |
| Lack of docility | 0.718 | 0.00 |
| Neophobia | 0.567 | 0.00 |
| Activity | 0.420 | 0.00 |
| **Glucocorticoids** |  |  |
| FCMs | 1.00 | 0.003 |
| **Innate immunity** |  |  |
| Neutrophils | 0.471 | 0.289 |
| Eosinophils | 0.245 | 0.150 |
| Monocytes | 0.075 | 0.046 |
| Basophils | 0.173 | 0.106 |
| Hemagglutination | 0.189 | 0.116 |
| Hemolysis | 0.135 | 0.083 |
| Outer model for adaptive immunity biomarkers | | |
| **Behavioural profile** |  |  |
| Lack of docility | 0.680 | 0.00 |
| Neophobia | 0.769 | 0.00 |
| Activity | 0.094 | 0.00 |
| **Glucocorticoids** |  |  |
| FCMs | 1.00 | 0.016 |
| **Adaptive immunity** |  |  |
| Gamma globulins | 0.338 | 0.238 |
| Lymphocytes | 0.568 | 0.401 |
| Anti-rabies antibody | 0.526 | 0.371 |
| Outer model for inflammatory immunity biomarkers | | |
| **Behavioural profile** |  |  |
| Lack of docility | 0.013 | 0.00 |
| Neophobia | 0.046 | 0.00 |
| Activity | 0.926 | 0.00 |
| **Glucocorticoids** |  |  |
| FCMs | 1.00 | 0.051 |
| **Inflammatory immunity** |  |  |
| Alpha-1 globulins | 0.876 | 0.554 |
| Alpha-2 globulins | 0.382 | 0.242 |
| Beta-globulins | 0.208 | 0.131 |
| Haptoglobin | 0.738 | 0.467 |
