## Supplementary material for "Co-variation between glucocorticoids, behaviour and immunity supports the pace-of-life syndrome hypothesis: an experimental approach": ESM S1, S2, S3, and S4: S3_ProcB_Carbillet.docx

^7 Equipe de Biologie médicale-Histologie, CREFRE, Inserm-UPS-ENVT, Toulouse, France^

^8 Inovie Vet, Laboratoire d'Analyses et Biologie Vétérinaires, Montpellier, France^

^9 Université de Toulouse, ENVT, INRAE, UMR IHAP, Toulouse, France.^

^10 ANSES, Nancy Laboratory for Rabies and Wildlife, Malzéville, France^

**Table S3.** Scores on the three axes (PC1, PC2, and PC3) of the principal component analysis (PCA) performed on the three behavioural parameters used in our study. Projections of all the variables on the different principal components and percentage of total inertia captured by each principal component. The first principal component (PC1) captured 51.7% of the total inertia and was much more informative than all lower order axes. Docility, Activity and Neophilia were all negatively correlated with PC1. Thus, this axis represented a gradient of behavioural profiles, with negative values indicating proactive behavioural profiles (high activity levels, neophobia, and lack of docility), and positive values indicating reactive behavioural profiles (low activity levels, neophilia, and docility).

| **Parameters** | **PC1** | **PC2** | **PC3** |
| --- | --- | --- | --- |
| Docility | -0.76 | -0.45 | 0.47 |
| Neophobia | -0.83 | -0.13 | -0.55 |
| Activity | -0.54 | 0.82 | 0.17 |
| Variance explained (%) | 51.7 | 29.8 | 18.4 |

**Fig. S3.** Projection of the three variables (in blue) considered and individuals (black points) on principal components 1 (x axis) and 2 (y-axis).

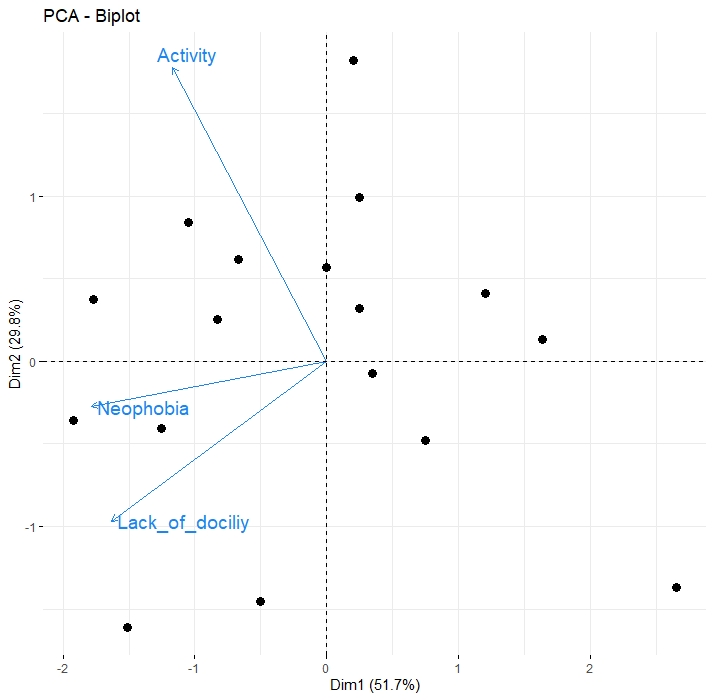
