## Supplementary material for "Co-variation between glucocorticoids, behaviour and immunity supports the pace-of-life syndrome hypothesis: an experimental approach": ESM S1, S2, S3, and S4: S4_ProcB_Carbillet.docx

^7 Equipe de Biologie médicale-Histologie, CREFRE, Inserm-UPS-ENVT, Toulouse, France^

^8 Inovie Vet, Laboratoire d'Analyses et Biologie Vétérinaires, Montpellier, France^

^9 Université de Toulouse, ENVT, INRAE, UMR IHAP, Toulouse, France.^

^10 ANSES, Nancy Laboratory for Rabies and Wildlife, Malzéville, France^

**Table S4.** Performance of the subset of candidate linear mixed-effect models within a ΔAICc < 2 fitted to investigate variation in baseline faecal glucocorticoids metabolite levels in the roe deer population of Gardouch. Model(s) in bold was/were used for estimation of parameters, and averaged when more than one model was considered after removing models that differed from a higher-ranking model by the addition of one or more parameters. These were rejected as uninformative, as recommended by Arnold (2010) and Richards (2008). Our set of candidate models was composed of all simpler models that included PC1 (behavioural profile ranging from proactive behavioural profiles to reactive behavioural profiles), age of individuals, period of sample collection and Julian date of sample collection. Individual identity and enclosure number were included as a random effect. AICc is the value of the corrected Akaike’s Information Criterion and K is the number of estimated parameters for each model. The ranking of the models is based on the differences in the values for ΔAICc and on the Akaike weights (AICw).

| **Models** | **K** | **AICc** | **ΔAICc** | **AICw** |
| --- | --- | --- | --- | --- |
| **PC1+Period** | **6** | **312.0** | **0.00** | **0.503** |
| PC1+Period+Julian date | 7 | 313.2 | 1.16 | 0.283 |
| PC1+Period+Age | 7 | 313.7 | 1.71 | 0.214 |
